# CRISPR-Cas9 *trans*-cleavage is hindered by a closed R-loop, an elongated spacer, and an inactive HNH domain

**DOI:** 10.1101/2025.04.17.649303

**Authors:** Roser Montagud-Martínez, Raúl Ruiz, Sara Baldanta, Rubén Delicado-Mateo, Guillermo Rodrigo

**Affiliations:** Institute for Integrative Systems Biology (I2SysBio), CSIC – University of Valencia, 46980 Paterna, Spain

**Keywords:** CRISPR-Cas systems, Dynamic behavior, Fluorogenic nucleic acids, Nuclease catalysis

## Abstract

Cas9 can process poly(T) single-stranded DNA molecules upon activation in an RNA-guided manner. Here, we uncover key structural determinants underlying this function. First, we show that open R-loops in the PAM-distal region favor *trans*-cleavage activity, which occur when targeting short double-stranded or single-stranded DNA molecules. Second, we show that elongated guide RNA spacers beyond the canonical 20 bases, even by a few bases, severely impairs this collateral activity. Third, although *trans*-cleavage is mediated by the RuvC domain, we show that a catalytically active HNH domain contributes to an efficient process. Structural analyses of domain rearrangements provide mechanistic insight. Together, these findings illustrate a fine modulation of Cas9 function.

## Letter

Several prokaryotes have adaptive immune systems encoded by clustered regularly interspaced short palindromic repeats (CRISPR) and genes expressing CRISPR-associated (Cas) proteins^1^. Due to their ever-growing functional versatility, these systems are being repurposed for a series of biotechnological applications, including gene editing, regulation, and detection^2^. Of note, the *trans*-cleavage activity upon nucleic acid targeting of both Cas12 and Cas13 has been instrumental to boost the field of CRISPR diagnostics^3,4^. Recently, the *trans*-cleavage activity of Cas9 has been disclosed over poly(T) single-stranded DNA (ssDNA) molecules^5^. This finding has been surprising because of a long-held opposite belief based on structural considerations. Such activity is apparently enhanced when the nuclease is guided in its native form with two RNA molecules, *i*.*e*., with a CRISPR RNA (crRNA) and a *trans*-activating crRNA (tracrRNA). Nevertheless, the sequence and structural determinants of this collateral catalytic activity are not fully appreciated.

To address this issue, we considered a set of nucleic acid targets in the form of short double-stranded DNA (dsDNA) containing the canonical protospacer adjacent motif (PAM), along with the corresponding crRNAs with a 20-22 nt spacer (more in detail, we used targets with a 20 bp stretch from the 5’ end to the PAM). We also considered the corresponding single guide RNAs (sgRNAs) for a comparative assessment. In this context, the R-loop that results from targeting with the CRISPR-Cas9 complex opens in the PAM-distal region^6^ (**Fig. 1A**). Using fluorogenic dsDNA molecules and a fluorogenic 18 nt poly(T) ssDNA probe, we scrutinized the targeting ability and *trans*-cleavage activity of the *Streptococcus pyogenes* Cas9 in each case (**Figs. 1B, 1C**). According to our results, both functions are not aligned, as *e*.*g*. in one case a high targeting ability led only to a moderate *trans*-cleavage activity with both the crRNA-tracrRNA and sgRNA (system #2), while in another case a low targeting ability resulted in a high *trans*-cleavage activity with the crRNA-tracrRNA (#4). We also observed that while dsDNA targeting is higher with either the crRNA-tracrRNA or sgRNA depending on the system (*e*.*g*., #1 *vs*. #4), ssDNA *trans*-cleavage is always higher or at least equal when Cas9 is guided in its native form. Some basal *trans*-cleavage activity (*i*.*e*., in absence of target DNA) was also noticed (**Fig. S1A**). These results hence confirm the new catalytic feature and indicate a sequence dependence on both the *cis*- and *trans*-actions.

**Figure 1.**
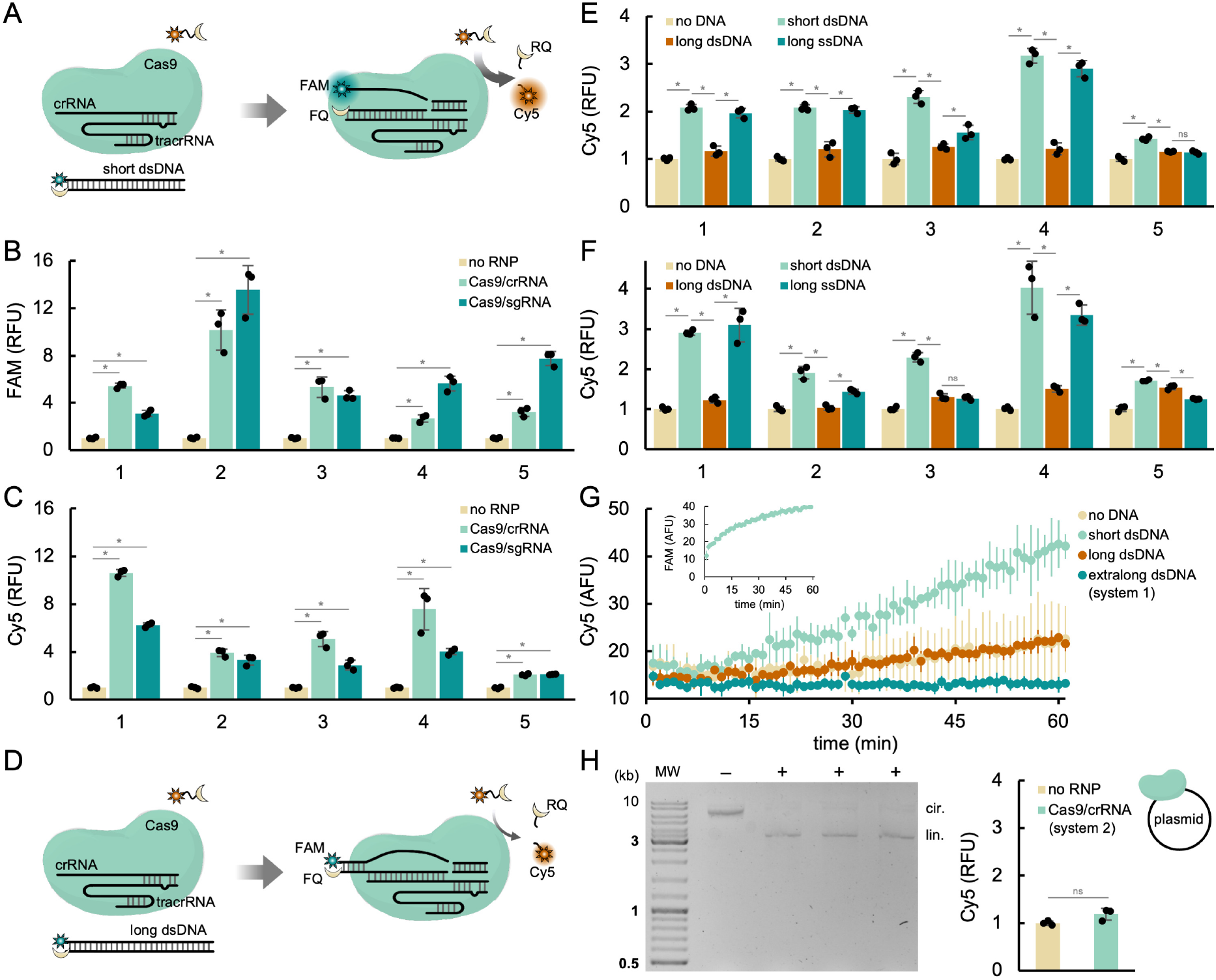
*Trans*-cleavage activity is hindered by a closed R-loop. A) Schematics of the CRISPR-Cas9 reaction used to analyze targeting ability (FAM signal) and *trans*-cleavage activity (Cy5 signal). Open R-loop in the PAM-distal region as a result of targeting a short dsDNA molecule. B) Targeting ability and C) *trans*-cleavage activity of five different systems. Cas9 guided by crRNA-tracrRNAs or sgRNAs with spacers of 20-22 nt. Fluorescence normalization by non-targeted state values. D) Schematics of the CRISPR-Cas9 reaction when using a long dsDNA target and the R-loop remains closed. E, F) *Trans*-cleavage activity with different DNA inputs: short dsDNA (open R-loop), long dsDNA (closed R-loop), and long ssDNA (no R-loop). Cas9 guided by crRNA-tracrRNAs in (E) and by sgRNAs in (F). Fluorescence normalization by DNA-free state values. G) Time-dependent *trans*-cleavage activity of system #1 with different DNA inputs (including an extralong dsDNA). In the inset, time-dependent targeting ability. H) *Cis*-cleavage of a plasmid (system #2) revealed by a gel electrophoretic assay and associated *trans*- cleavage activity on the right. GeneRuler DNA ladder mix (100-10000 bp). Cas9 guided by a crRNA-tracrRNA in (G), (H). Represented data in the bar plots correspond to means plus/minus standard deviations, along with the individual points (*n* = 3). Statistical significance assessed by two-tailed Welch’s *t*-tests (^*^*P* < 0.05, ^ns^non-significant change). RNP, ribonucleoprotein. FAM, 6-Carboxyfluorescein. Cy5, Cyanine 5. FQ, Iowa Black FQ. RQ, Iowa Black RQ. RFU, relative fluorescent units. AFU, absolute fluorescent units.

To distill a clear structural determinant, we next considered longer dsDNA molecules to target (35-36 bp stretch from the 5’ end to the PAM). In this case, the resulting R-loop remains closed in the PAM-distal region if using a crRNA or sgRNA with a 20-22 nt spacer (**Fig. 1D**). Intriguingly, we found a systematic substantial reduction in *trans*-cleavage activity with respect to the use of short dsDNA inputs (**Figs. 1E, 1F**). In the case of system #4, *e*.*g*., the reduction was 90% by guiding Cas9 with the crRNA-tracrRNA and 81% with the sgRNA. Here, *cis*-cleavage was confirmed by gel electrophoresis (**Fig. S1B**). We then found that by targeting long ssDNA molecules with crRNA-tracrRNAs (in a PAM-independent way)^7^ the *trans*-cleavage activity was almost recovered, except in one case in which this process was not efficient (#5). Similar results were obtained with sgRNAs. Using a fluorogenic 13 nt poly(T) ssDNA probe, we also observed a decline in its processing when the R-loop was closed (**Fig. S1C**). Yet, it was noteworthy the higher *trans*-cleavage activity in the case of systems #1 and #4 with long ssDNA inputs. By contrast, no apparent *trans*-cleavage activity was observed with a fluorogenic 18 nt poly(A) ssDNA probe (**Fig. S1D**). Thus, our data entail that a closed R-loop hinders the collateral catalytic activity of Cas9.

To further study the dynamics of the response, we monitored over time the fluorescence signals associated with dsDNA targeting and subsequent ssDNA *trans*- cleavage (for system #1 with the crRNA-tracrRNA). *Cis*-targeting was fast, with a mean reaction time of ~7 min, but a delay of ~10 min was found in *trans*-cleavage together with a slower reaction rate (**Fig. 1G**). Of note, ssDNA collateral catalysis was completely abolished, even the basal activity of the ribonucleoprotein, using an extralong dsDNA target (49 bp stretch from the 5’ end to the PAM), in tune with the formation of a more stable closed R-loop. Indeed, this behavior was reproduced when targeting a plasmid (**Fig. 1H**). This suggests that this feature might play a limited role in the natural *in vivo* context.

Subsequently, we reasoned that the collateral activity of Cas9 could be modulated in a quantitative manner through R-loop reconfigurations. Indeed, we observed that the longer the DNA stretch from the 5’ end to the PAM, the lower the *trans*-cleavage activity (**Fig. 2A**; Cas9 guided by crRNA-tracrRNAs, relative to system #1). More in detail, such activity correlated with the free energy of the dsDNA target in the PAM-distal region calculated from an empirical thermodynamic model^8^. Then, we analyzed the effect of simultaneous crRNA spacer elongations to maintain an open R-loop in the PAM-distal region. Surprisingly, we found that poly(T) probe processing was disrupted when using these longer crRNAs (**Fig. 2B**; see also **Figs. S2A, S2B**). To shed light, we created a series of crRNAs with strict spacer lengths (including the GR nucleotides in the 5’ end required for *in vitro* transcription with a T7 RNA polymerase). Elongating the crRNA spacer roughly maintained the DNA binding capacity (**Fig. 2C**); a feature that has already been exploited for nucleic acid detection^9^. However, *trans*-cleavage activity decreased with spacer length following a nearly linear trend (**Fig. 2D**). In particular, the reduction was 49% with a spacer of strict 22 nt and 79% with 24 nt. We also observed a slight increase in *trans*-cleavage activity with respect to the canonical case using a crRNA with 19 nt spacer. This agrees with previous results showing reduction in on-target cleavage activity by modifying the 5’ end of the sgRNA^10^. Therefore, our data demonstrate that spacer length is a critical variable to finely control the collateral activity of Cas9.

**Figure 2.**
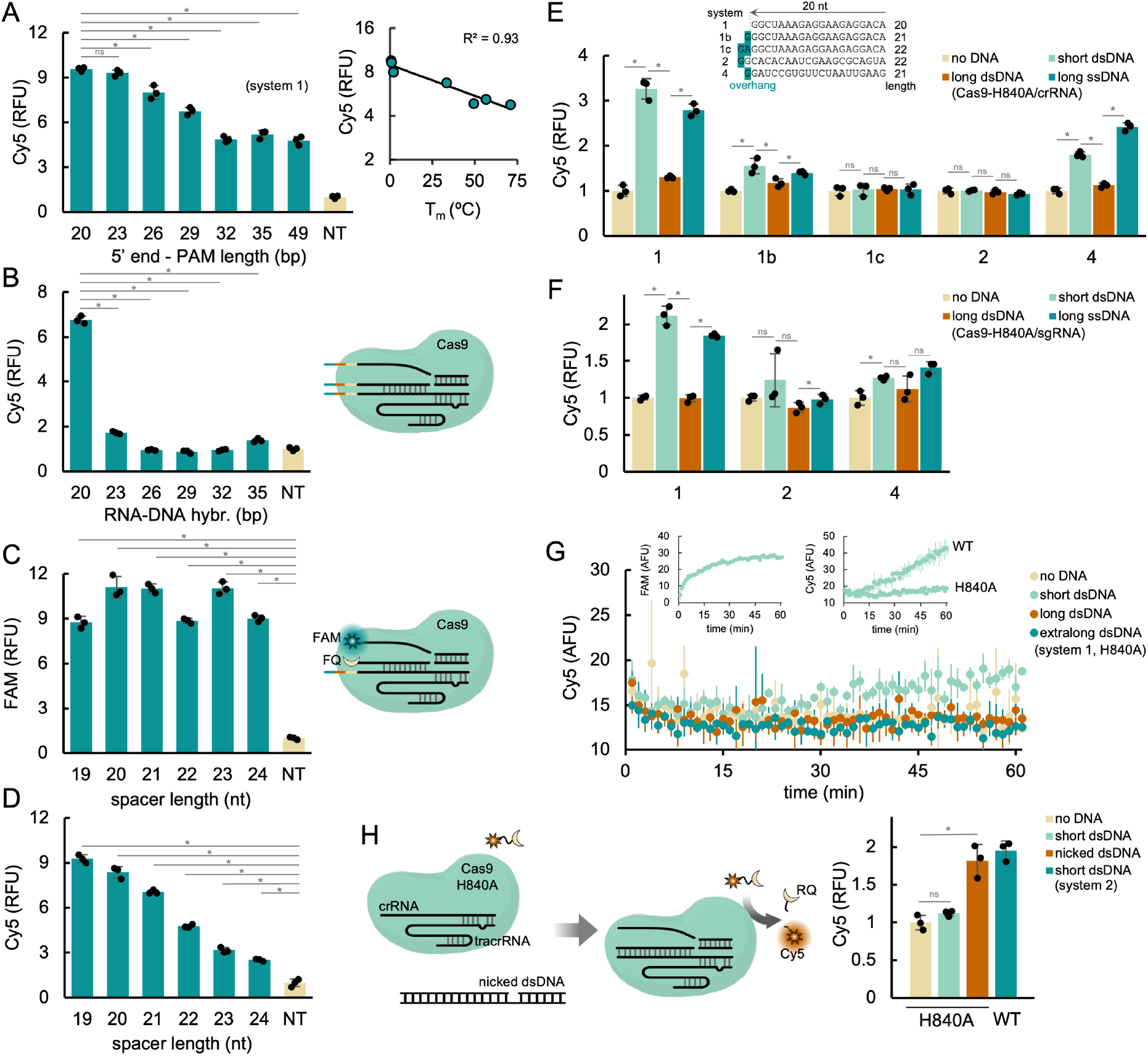
*Trans*-cleavage activity is hindered by an elongated spacer and an inactive HNH domain. A, B) *Trans*-cleavage activity of system #1 when varying the R-loop configuration. Extension of the stretch from the 5’ end to the PAM in the dsDNA input keeping fixed the crRNA with a 20 nt spacer in (A). On the right, correlation between *trans*- cleavage activity and melting temperature of the PAM-distal region of the dsDNA input. Extension of the RNA-DNA hybridization region in (B), varying simultaneously the length of the crRNA spacer and the stretch from the 5’ end to the PAM in the dsDNA input. On the right, scheme of the system. C) Targeting ability and D) *trans*-cleavage activity of system #1 when varying the length of the crRNA spacer keeping fixed the DNA input (short dsDNA, open R-loop). On the right, scheme of the system. Fluorescence normalization by non-targeted state values (NT, non-targeting crRNA). Cas9 guided by crRNA-tracrRNAs in (A) # (D). E, F) *Trans*-cleavage activity of various systems with HNH-mutant Cas9 and different DNA inputs: short dsDNA (open R-loop), long dsDNA (closed R-loop), and long ssDNA (no R-loop). The inset shows the respective crRNA spacers of 20-22 nt. Cas9 guided by crRNA-tracrRNAs in (E) and by sgRNAs in (F). Fluorescence normalization by DNA-free state values. G) Time-dependent *trans*-cleavage activity of system #1 with HNH-mutant Cas9 and different DNA inputs (including an extralong dsDNA). In the insets, time-dependent targeting ability (left) and *trans*-cleavage activity comparative against wild-type Cas9 (right). H) Schematics of the CRISPR-Cas9 reaction used to analyze the effect of a nicked dsDNA input and associated *trans*-cleavage activity on the right. Cas9 guided by a crRNA-tracrRNA in (G), (H). Represented data in the bar plots correspond to means plus/minus standard deviations, along with the individual points (*n* = 3). Statistical significance assessed by two-tailed Welch’s *t*-tests (^*^*P* < 0.05, ^ns^non-significant change). FAM, 6-Carboxyfluorescein. Cy5, Cyanine 5. FQ, Iowa Black FQ. RQ, Iowa Black RQ. RFU, relative fluorescent units. AFU, absolute fluorescent units.

Furthermore, we investigated the impact of perturbing the nuclease function. Cas9 is a large multidomain protein that contains two endonuclease domains, RuvC and HNH, along with a PAM-recognizing domain^11^. According to Chen *et al*.^5^, RuvC is the domain responsible for *trans*-cleavage. Using the HNH-mutant Cas9 (working like a nickase)^12^ and crRNA-tracrRNAs, we here found that such a domain also plays a role in *trans*-cleavage

(**Fig. 2E**; see also the null functionality of the RuvC-mutant Cas9 in **Figs. S2C, S2D**). Absolute fluorescence levels associated with *trans*-cleavage were substantially lower. The effect of the HNH domain was further evidenced by using crRNAs of different length. While the wild-type nuclease was still able to process the poly(T) probe with significant efficacy when using a crRNA with a 22 nt spacer, the HNH-mutant Cas9 failed to do so (*e*.*g*., #1 *vs*. #1c), only displaying collateral activity if guided by crRNAs with a 21 nt spacer at most. Moreover, this mutant tolerated less well the use of sgRNAs (**Fig. 2F**). The time-dependent fluorescence signals associated with the *cis*- and *trans*-actions (for system #1) were also monitored (**Fig. 2G**). Interestingly, employing a nicked dsDNA input in the targeted strand and a crRNA with a 22 nt spacer (for system #2), we found that the HNH-mutant Cas9 displayed similar collateral activity to the wild-type nuclease (**Fig. 2H**). Arguably, a break in the targeted strand provides to Cas9 the required structural flexibility to efficiently activate the RuvC domain^13^.

To gain mechanistic insight, we analyzed a set of three-dimensional structures deposited in the Protein Data Bank (PDB). A capping of the 5’ end of the sgRNA, with contacts with the RuvC domain, was observed in a structure corresponding to a CRISPR-Cas9 complex bound to a short dsDNA molecule in which a strict 20 nt spacer follows and the R-loop presents an open configuration^14^ (**Fig. 3A**). The capping seems to hold in a closed R-loop scenario, which results from targeting a long dsDNA molecule, if the spacer still has the canonical length^15^ (**Fig. 3B**). However, the analysis of the structure of a complex in which the sgRNA has an elongated spacer (GG overhang in its 5’ end) revealed the exposition to the solvent of that tail^16^ (**Fig. 3C**). In addition, we observed changes in the conformation of the central channel formed between the recognition and nuclease lobes where the DNA substrate is accommodated. If the R-loop is closed in the PAM-distal region, the space between the REC3 and RuvC domains narrows (**Figs. S3A, S3B**). Accordingly, the RuvC active site might become less solvent-accessible, leading to lower processing ability of outer ssDNA molecules. Moreover, it is known that Cas9 adopts a more compact conformation around the central channel once it is primed for catalysis, especially because the HNH domain docks at the scissile phosphate of the targeted strand and the recognition lobe reorients to stabilize DNA binding^14^. Despite, the RuvC domain remains relatively stable during activation. Yet, we noticed different configurations of this domain in the **Fig. 3A** (catalytic state, capped 5’ end) and **Fig. 3C** structures (checkpoint state, exposed 5’ end). In particular, the intradomain distance between G1011 (a potential RNA-interacting residue) and L1052 (a residue in the groove where the non-targeted strand is accommodated) changes from 32.9 to 25.2 Å. In this regard, the capping of the 5’ end of the sgRNA might contribute to have an optimal configuration of the RuvC domain for an efficient cleavage activity (both in *cis* and *trans*). This information provides, at least in part, a structural basis for the differential collateral activity here reported, stressing the high plasticity of the Cas proteins.

**Figure 3.**
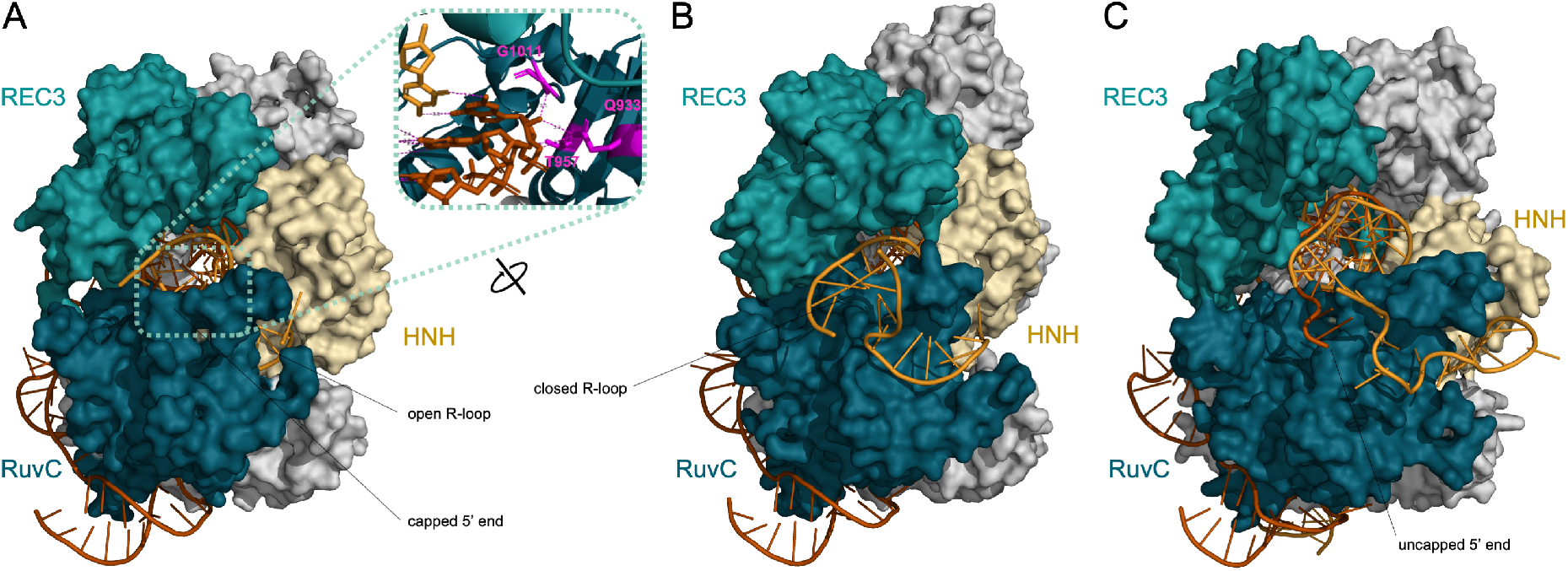
Conformational states underlying *trans*-cleavage activity. A) Three-dimensional structure of a CRISPR-Cas9 complex bound to a short dsDNA molecule in which Cas9 is catalytically active, the sgRNA has a strict 20 nt spacer, and the R-loop presents an open conformation (PDB #5F9R). In the zoom-in inset, interactions of the 5’ end of the sgRNA with Cas9, which contribute to the sgRNA capping (RNA-interacting residues with nucleotide at position 20 shown in magenta). B) Three-dimensional structure of a CRISPR-Cas9 complex bound to a long dsDNA molecule in which Cas9 is catalytically active, the sgRNA has a 20 nt spacer (GG overhang in the 5’ end), and the R-loop presents an open conformation (PDB #7S4X). C) Three-dimensional structure of a CRISPR-Cas9 complex bound to a long dsDNA molecule in the checkpoint state in which Cas9 is catalytically dead (dCas9), the sgRNA has a 22 nt spacer (GG overhang in the 5’ end), and the R-loop presents a closed conformation (PDB #5Y36). The HNH, RuvC, and REC3 domains are colored in pale yellow, blue, and turquoise, respectively, while the rest of the protein is colored in white. The sgRNA is colored in orange and the DNA strands in yellow. For representation purposes, the target DNA molecule was trimmed after the PAM. Rendering obtained with PyMOL^17^.

Conclusively, this work provides overlooked structural determinants of the CRISPR-Cas9 *trans*-cleavage activity. Closed R-loops in the PAM-distal region and elongated spacers beyond the canonical 20 nt are detrimental. Moreover, a catalytically active HNH domain favors *trans*-cleavage activity, despite it is accomplished by the RuvC domain, which aligns with the fact that these two nuclease domains act in concert^13^. It seems that *S. pyogenes* Cas9 evolved to prioritize precise on-target dsDNA cleavage, but these findings expand our view to consider as well its restricted collateral activity. We expect they will inspire developments in biological engineering and diagnostics. Further work is needed to define sequence determinants, resolve atomic structures in additional functional states, and explore *trans*-cleavage behavior across nuclease variants. As our understanding of CRISPR-Cas systems grows, so does our capacity to translate them into tools that address societal challenges^2^.

## Acknowledgements

Work supported by the Spanish Ministry of Science, Innovation, and Universities and AEI/10.13039/501100011033 (PDC2022-133941-I00 and PID2021-127671NB-I00, co-financed by the European Union NextGenerationEU/PRTR and European Regional Development Fund) and the Valencia Regional Government (CIPROM/2022/21). SB acknowledges a Juan de la Cierva contract from the Spanish Ministry of Science, Innovation, and Universities (JDC2023-052427-I) and RDM a predoctoral contract from the Valencia Regional Government (CIACIF/2023/119).

## Authors’ contribution

GR designed the research. RMM performed the experiments under the supervision of GR and supported by RR, SB, and RDM. All authors analyzed the data. GR wrote the manuscript. All authors revised the final manuscript.

## ONLINE METHODS

### Molecular elements

Target DNA molecules were chemically synthesized and purified by IDT. They were labelled with the green fluorophore 6-Carboxyfluorescein (FAM) and the dark quencher Iowa Black FQ in their PAM-distal ends, so that we could monitor the targeting ability. The IDT OligoAnalyzer tool was used to calculate the melting temperatures of the PAM-distal regions of some target molecules specifying a DNA concentration of 40 nM and a Mg^2+^ concentration of 12.5 mM. Poly(T) ssDNA probes of 18 nt and 13 nt, as well as a poly(A) ssDNA probe of 18 nt, were chemically synthesized and purified by IDT. They were labelled with the far-red fluorophore Cyanine 5 (Cy5) and the dark quencher Iowa Black RQ in their ends, so that we could monitor the *trans*-cleavage activity. Three versions of *S. pyogenes* Cas9 (IDT) were used: the wild-type nuclease, the H840A nickase (HNH mutant), and the D10A nickase (RuvC mutant). In addition, the sgRNAs, crRNAs, and tracrRNA were produced *in vitro* with the TranscriptAid T7 high yield transcription kit (Thermo) from DNA templates. They were purified using the RNA clean and concentrator column (Zymo) and quantified in a spectrophotometer (NanoDrop, Thermo). Sequences provided in **Table S1**.

### CRISPR-Cas9 reactions

Reactions were performed in 1x Tris/Acetate/EDTA (TAE) buffer pH 8.5 (Invitrogen), 0.05% Tween 20, and 12.5 mM MgCl_2_ at a final volume of 20 µL. 200 nM CRISPR-Cas9 ribonucleoprotein, previously assembled at room temperature for 30 min, was mixed with 40 nM target DNA (in the form of dsDNA or ssDNA) and 100 nM ssDNA probe. Reactions were incubated at 37 ºC for 1 h in tubes in a thermocycler (Mastercycler, Eppendorf). For time-dependent activity analyses, reactions were directly incubated at 37 ºC for 1 h in a microplate in the fluorometer.

### Fluorescence quantification

CRISPR-Cas9 reaction volumes were loaded in a black 384-well microplate with clear bottom (Falcon), which was then placed in a fluorometer (CLARIOstar Plus, BMG) to measure green and far-red fluorescence. For FAM, excitation was at 490/8 nm and emission at 526/12 nm (green); for Cy5, excitation was at 640/8 nm and emission at 675/12 nm (far-red). The fluorescence values of the reaction buffer were subtracted to correct the signals. Absolute or relative fluorescence values were represented in the figures.

### Gel electrophoresis

Nucleic acid cleavage by Cas9 was confirmed by agarose gel electrophoresis. For that, CRISPR-Cas9 reactions were first treated with 10 µg proteinase K (Ambion), incubating at 50 ºC for 1 h. The digestion was inactivated at 95 ºC for 15 min. Subsequently, 1 U RNase H (Ambion) and 2 µg RNase A (Invitrogen) were added, incubating at 37 ºC for 1 h. Samples were then loaded on a 1% or 3% agarose gel prepared with 0.5x Tris/Borate/EDTA (TBE) buffer, which was run for 45 min at room temperature (110 V). Gels were stained using GreenSafe (NZYtech). The GeneRuler DNA ladder mix (100-10000 bp, Thermo) and ultra-low range DNA ladder (10-300 bp, Thermo) were used as markers.

### Structural models

Three different structures of a CRISPR-Cas9 complex bound to a target DNA molecule were analyzed: *i*) the crystal structure of a complex bound to a short dsDNA molecule in which the sgRNA has a strict 20 nt spacer (PDB #5F9R), *ii*) the cryogenic electron microscopy (cryo-EM) structure of a complex bound to a long DNA molecule in which the sgRNA has a strict 20 nt spacer (PDB #7S4X), and *iii*) the cryo-EM structure of a complex bound to a long dsDNA molecule in which the sgRNA has a 22 nt spacer (GG overhang in the 5’ end; PDB #5Y36). These structures were visualized and aligned using PyMOL. In the structures #5F9R and #7S4X, Cas9 was catalytically active; while in the structure #5Y36, Cas9 was catalytically dead (harboring the D10A and H840A mutations).

## SUPPLEMENTARY FIGURE/TABLE LEGENDS

**Figure S1. Extended data related to the effect of the R-loop configuration on *trans*- cleavage function**. A) Basal *trans*-cleavage activity of Cas9 (no DNA target) when there is no guide RNA and when it is guided by crRNA-tracrRNAs or sgRNAs (with spacers of 20-22 nt). B) Gel electrophoretic assay to show *cis*-cleavage when targeting long dsDNA molecules. GeneRuler ultra-low range DNA ladder (10-300 bp). C, D) *Trans*-cleavage activity with different DNA inputs: short dsDNA, long dsDNA, and long ssDNA. Cas9 guided by crRNA-tracrRNAs. Poly(T) ssDNA probe of 13 nt in (C) and poly(A) ssDNA probe of 18 nt in (D). AFU, absolute fluorescence units.

**Figure S2. Extended data related to the effect of spacer length and HNH domain activity on *trans*-cleavage function**. A) Targeting ability when varying the length of the crRNA spacer keeping fixed the DNA input (long dsDNA, system #1). Fluorescence normalization by the value obtained with a 20 nt spacer. B) *Trans*-cleavage activity with crRNA spacers of different length (long dsDNA, system #1). 29* indicates a spacer elongated in 9 uracils. C, D) *Trans*-cleavage activity with RuvC-mutant Cas9 and different DNA inputs: short dsDNA, long dsDNA, and long ssDNA. Cas9 guided by crRNA-tracrRNAs in (C) and by sgRNAs in (D). Fluorescence normalization by DNA-free state values. RFU, relative fluorescent units. AFU, absolute fluorescence units.

**Figure S3. Extended data related to the structural analysis of *trans*-cleavage function** A, B) Alternative view of Cas9 showing the relative position of the RuvC and REC3 domains that interact with the PAM-distal region. Representation from PDB #5F9R in A) and from PDB #7S4X in B). An arrow points to the space between the REC3 and RuvC domains. HNH is shown in pale yellow, RuvC in blue, REC3 in turquoise, sgRNA in orange, targeted DNA in yellow.

**Table S1**. Sequences of the different nucleic acids used in this work. In the target DNA molecules, PAMs are marked in bold.

